# The *Pseudomonas aeruginosa* Wsp pathway undergoes positive evolutionary selection during chronic infection

**DOI:** 10.1101/456186

**Authors:** Erin S. Gloag, Christopher W. Marshall, Daniel Snyder, Gina R. Lewin, Jacob S. Harris, Sarah B. Chaney, Marvin Whiteley, Vaughn S. Cooper, Daniel J. Wozniak

## Author Contributions

ESG and CWM performed the experimental work. JSH performed the colony PCR. SBC infected and sampled the porcine burn wounds. DS generated the sequence library. MW and GRL quantified the strain frequency in the wounds. ESG, CWM, MW, VSC, and DJW conceptualized the project and wrote the manuscript.

## Introductory paragraph

Pathogens experience pressure in an infection to adapt, with selection favoring mutants that persist. *Pseudomonas aeruginosa* commonly adapts by evolving mutants with hyper-biofilm production that evade clearance. Despite our understanding of the adaptive phenotypes, studying their emergence and dynamics in an infection has proven challenging. Here we used a porcine full-thickness burn wound model of chronic infection to study how mixed strains of *P. aeruginosa* adaptively evolve. Wounds were infected with six *P. aeruginosa* strains, including the model PA14 strain (PA14-1), and biopsies taken at 3, 14, and 28 days post-infection. Rugose small-colony variants (RSCVs) were detected at 3-d and persisted, with the majority evolved from PA14-1. Whole genome sequencing of PA14-1 RSCVs revealed driver mutations exclusively in the *wsp* pathway. RSCVs also acquired CRISPR-Cas adaptive immunity to prophages isolated from the *P. aeruginosa* wound isolate (B23-2) present in the inoculum. The rapid rise of RSCVs to detectable frequencies is evidence of positive selection of the Wsp chemosensory system and suggests that RSCVs may arise earlier in an infection than originally appreciated, to facilitate infection. Given the prevalence of RSCVs in chronic infections, we predict that RSCVs may be a common, early adaptation during infections.

Chronic infections are those that persist despite extensive treatment. These persistent infections are often attributed to difficult to eradicate biofilms, which are communities of adhered microorganisms encased in an extracellular polymeric substance (EPS)^1^^,^^2^. Complicating chronic infections is the high likelihood that bacterial populations adaptively evolve, producing persistent phenotypes with increased fitness.

One of the most understood bacterial adaptive responses to chronic infection is that of *Pseudomonas aeruginosa* to the cystic fibrosis (CF) lung^3^. CF patients exhibit mucus accumulation, where *P. aeruginosa* biofilms commonly colonize the mucus lining and establish persistent pulmonary infections^4^^-^^6^. Evolved phenotypic variants of this organism are routinely isolated from CF patient sputum samples, and several of these variants are often associated with worsening patient prognosis^7^. Of particular interest are the rugose small-colony variants (RSCVs), which are isolated from up to 50% of *P. aeruginosa*-positive CF sputum samples^8^^,^^9^. When isolated, their frequencies range drastically between 0.1 − 100%^9^^,^^10^. In contrast there is little-to-no reports of the frequency of *P. aeruginosa* adapted variants from other chronic infections, such as chronic wound infections.

Common to RSCVs are mutations in pathways that lead to elevated cyclic diguanylate monophosphate (c-di-GMP)^7^. C-di-GMP is a messenger molecule that signals the transition from planktonic to biofilm lifestyle in many bacteria^11^. In *P. aeruginosa*, increased c-di-GMP, among many responses, leads to overproduction of exopolysaccharides, Psl and Pel, and matrix proteins^12^^,^^13^. As a result, RSCVs have hyper-biofilm phenotypes^13^^,^^14^, increased tolerance to antimicrobials^10^, and enhanced resistance to immunity^15^^,^^16^. *P. aeruginosa* RSCVs also evolve from *in vitro* grown biofilms^17^^,^^18^, suggesting that there is strong selection for ecological diversification in both *in vivo* and *in vitro* biofilms.

RSCVs with driver mutations in the Wsp pathway, particularly *wspF,* are commonly isolated from CF sputum and *in vitro* biofilms^12^^,^^19^. The Wsp (wrinkly spreader) pathway is a chemosensory system that regulates c-di-GMP in response to surface sensing. Upon detecting a surface, the methyl-accepting chemotaxis protein (MCP), WspA, is methylated by the methyltransferase, WspC. WspA then interacts with the histidine kinase, WspE, which phosphorylates the diguanylate cyclase (DGC) WspR, resulting in c-di-GMP synthesis^20^^-^^23^. The methylesterase, WspF, de-methylates WspA, re-setting the system. Therefore, in *wspF* loss of function mutants, WspA remains methylated and the Wsp pathway continually activated, leading to overproduction of c-di-GMP^24^.

Despite our understanding of the divergent phenotypes of evolved variants, studying their emergence and the selective pressures driving their evolution *in vivo* is challenging, as there are few chronic infection models that mimic what is observed clinically. To address this challenge, we used a porcine full-thickness thermal injury wound model, which closely reflects human clinical chronic wounds^25^^,^^26^. Furthermore, chronic infection models typically only address the adaptive traits of a single founding clone, but susceptible individuals are constantly exposed to different strains of opportunistic pathogens. Here we used the porcine wound model in a *P. aeruginosa* mixed strain infection to understand which strains become prevalent and how they undergo genetic and phenotypic diversification.

## Results

### *P. aeruginosa* strains PA14-1 and PAO1-B11 become dominant in a *P. aeruginosa* mixed-strain chronic burn wound infection

To determine relative fitness of different *P. aeruginosa* strains and if the population evolves in a chronic infection, we infected porcine full-thickness burn wounds with an inoculum consisting of approximately equal numbers of 6 different *P. aeruginosa* strains. Wounds were infected with 2 model strains (PA14-1 and PAO1-B11), 3 clinical isolates (B23-2, CF18-1 and S54485-1), and a water isolate (MSH10-2) (Fig 1A, Table S1). Each strain had a unique nucleotide barcode introduced at the neutral *Tn7* site (Table S1). These strains share similar metabolic kinetics (Fig S1A, B) and biofilm formation capacity (Fig S1C). Biopsies were taken 3-, 14-, and 28-d post-infection for bacterial quantification (Fig S2).

**Figure 1:**
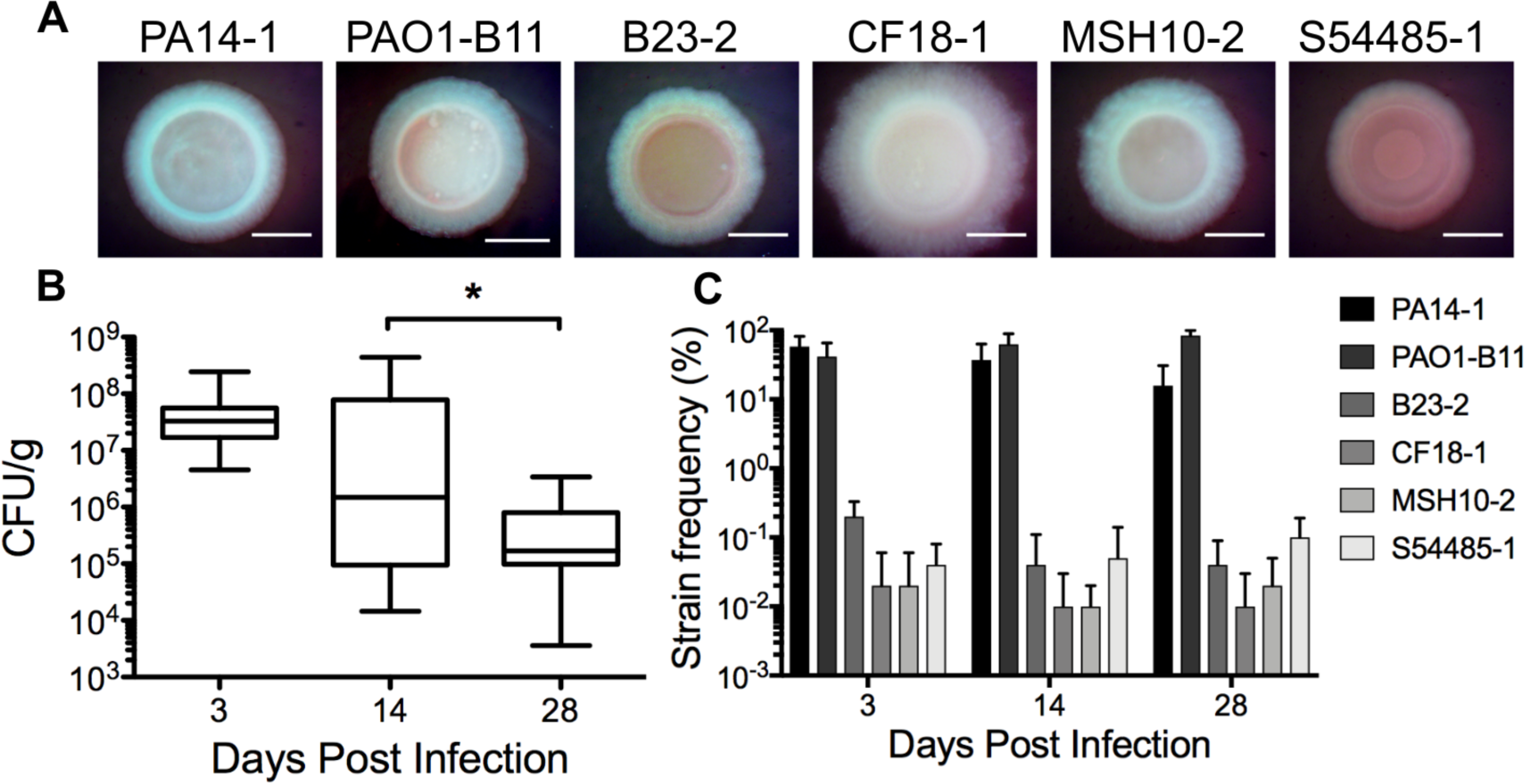
*P. aeruginosa* burden in a mixed-strain chronic burn wound infection. (**A)** Colony morphology on VBMM of the 6 *P. aeruginosa* strains used to infect porcine burn wounds. Scale bar 2mm. (**B)** Biopsies were taken from wounds at 3, 14 and 28-d. Biopsies were homogenized and plated for CFU/g. Each biopsy was plated in triplicate, with a minimum of 4 biopsies taken from each wound. * p-value <0.05. (**C)** Genomic DNA was isolated from homogenized biopsies and the strain specific barcodes at the *Tn7* site sequenced. The proportion of strain barcodes was expressed as a percentage of the total sequence reads to determine the relative frequency of each strain during the infection. A minimum of 4 biopsies from each wound was sequenced.

Colony forming units (CFU) revealed a high bacterial burden at both 3-d and 14-d post infection. However, the bacterial levels from 14-d biopsies were more variable across replicates (Fig 1B), suggesting wounds had begun clearing the infection. Wounds remained colonized at approximately 10^5^ bacteria up to 28-d (Fig 1B). To quantify the proportion of each strain across the sampled timepoints, genomic DNA was isolated from biopsy tissue and sequenced. As early as 3-d post infection, PA14-1 and PAO1-B11 were the predominant strains in the infection, outcompeting the 4 other strains present (Fig 1C).

### *P. aeruginosa* evolves RSCVs during porcine chronic burn wound infections

To determine if adapted *P. aeruginosa* variants emerged, we used colony morphologies as an indicator. Homogenized biopsies were grown on Vogel-Bonner minimal media supplemented with Congo red and brilliant blue (VBMM). RSCVs, defined by a matte, rugose colony morphology that stained intensely by the two dyes, indicating exopolysaccharide overproduction, were isolated from all three timepoints (Fig 2A). Two RSCV sub-populations were observed; one that had a pink, rugose phenotype and a second that had an orange, textured phenotype (Fig 2A).

**Figure 2:**
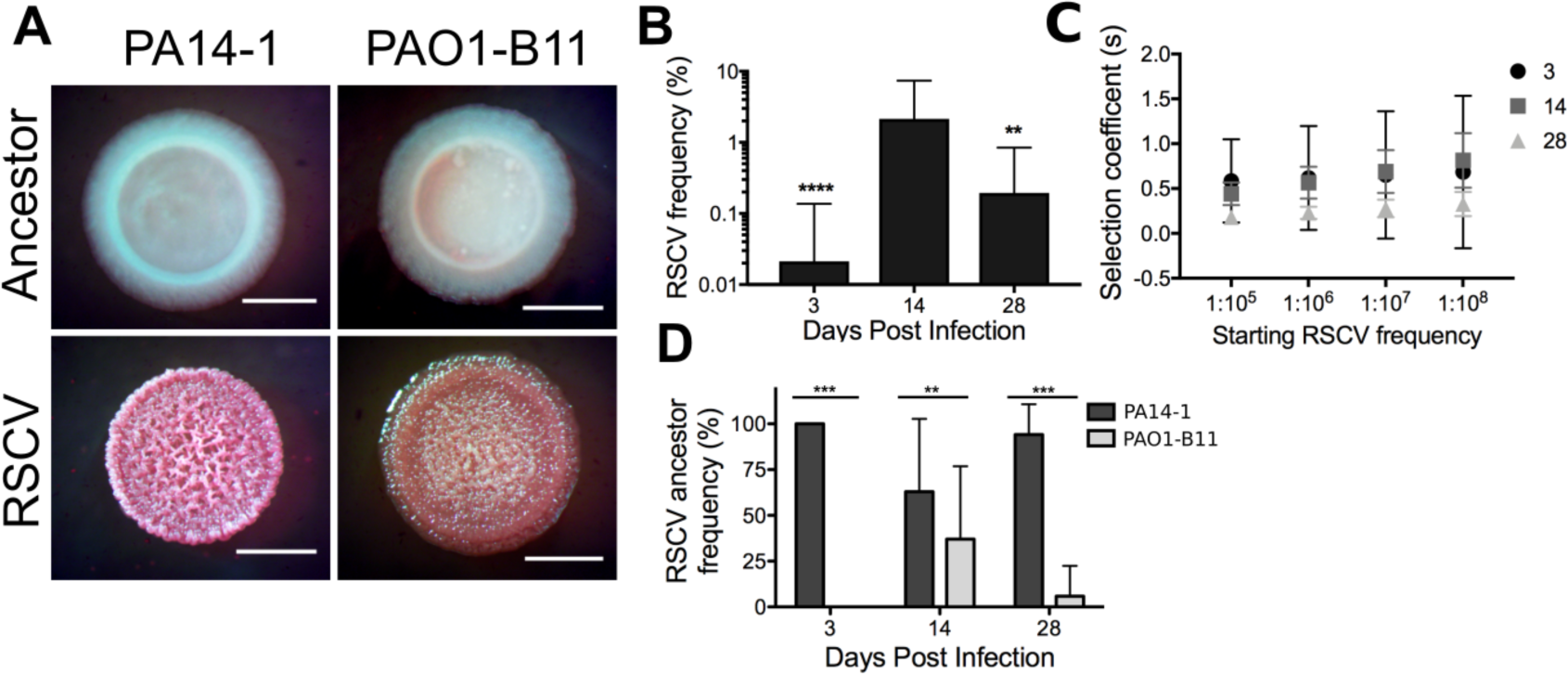
*P. aeruginosa* RSCVs isolated from porcine chronic burn wound infections. (**A)** Representative colony morphologies of RSCVs isolated from homogenized porcine burn wound tissue. RSCVs were plated on VBMM and the colony morphology was compared to the ancestor strain (labelled). Scale bar 2mm. (**B)** Frequencies of RSCVs isolated at each timepoint, expressed as a percentage of the total *P. aeruginosa* population. Data presented as mean ± SD. ** p-value <0.01, **** p-value <0.0001 compared to 14-d. (**C)** Selection coefficient (*s*) of RSCVs in the wound determined at each timepoint across a range of starting frequencies at t=0. (**D)** Frequency of the ancestor strain that the RSCVs evolved from, expressed as a percentage of the total RSCV sub-population. Data presented as mean ± SD. ** p-value <0.01, *** p-value <0.001.

The RSCV abundance in the wounds was quantified by expressing their frequency as a percentage of the total *P. aeruginosa* burden. The RSCV frequency was low in the wounds on 3-d (0.02 ± 0.12%) before peaking at 14-d, with a frequency of approximately 2% (2.15 ± 5.25%) (Fig 2B). On 28-d, the RSCV frequency decreased to 0.19 ± 0.65% of the total bacterial burden (Fig 1B, 2B). To determine if RSCVs experienced selective pressure in the wound, we calculated the selection coefficient, relative to the inoculum, at each timepoint across a range of possible starting frequencies according to equation (1). RSCVs showed significant positive selection across all time points with *s* >0.1 (Fig 2C). Selection of RSCVs was similar across all possible ranges on 3-d, while on 14-d and 28-d there was a gradual increase in selection coefficients. Despite the RSCV frequency decreasing on 28-d (Fig 2B), the RSCVs still showed significant positive selection in order to rise from undetectable frequencies to nearly 0.2% at 28-d (Fig 2C). This indicates that across all time points, RSCVs are adaptive.

RSCVs only evolved from the model strains PA14-1 and PAO1-B11 (Fig 2D). These also corresponded to the two RSCV phenotypes that were observed, with the pink RSCVs evolving from PA14-1, and the orange from PAO1-B11 (Fig 2A). RSCVs derived from PA14-1 were isolated across all timepoints and PAO1-B11 evolved RSCVs isolated from 14-d and 28-d. At the later two timepoints, PA14-1 RSCVs remained the predominant sub-population (Fig 2D).

### Whole genome sequencing reveals that PA14-1 RSCVs contain driver mutations exclusively within the *wsp* pathway

As PA14-1 RSCVs were the predominate evolved phenotype, we focused on this sub-population for the remainder of the study. A description of the PAO1-B11 variants will be communicated elsewhere. Whole genome sequencing was performed on 27 randomly selected PA14-1 RSCVs to identify the mutation(s) accounting for the RSCV phenotype.

We identified putative driver mutations exclusively in the *wsp* cluster, specifically, small deletions in *wspA* and *wspF.* (Table 1). We identified an in-frame 42bp deletion (Δ285 − 298aa) in *wspA* (*wspA* Δ285-298), and a frame-shift 5bp deletion (Δ461 − 465bp) in *wspF* (*wspF* V154fs) (Table 1). RSCVs with the *wspA* mutation were predominant across all timepoints, with *wspF* mutants only identified on 14-d. Using RSCV-2 as a representative *wspA* mutant, the variant *wspA* was replaced on the genome by a wildtype copy. This resulted in the RSCV colony phenotype reverting to wildtype (Fig 3), demonstrating that the *wspA* Δ285-298 mutation is responsible for the RSCV phenotype.

**Table 1:**
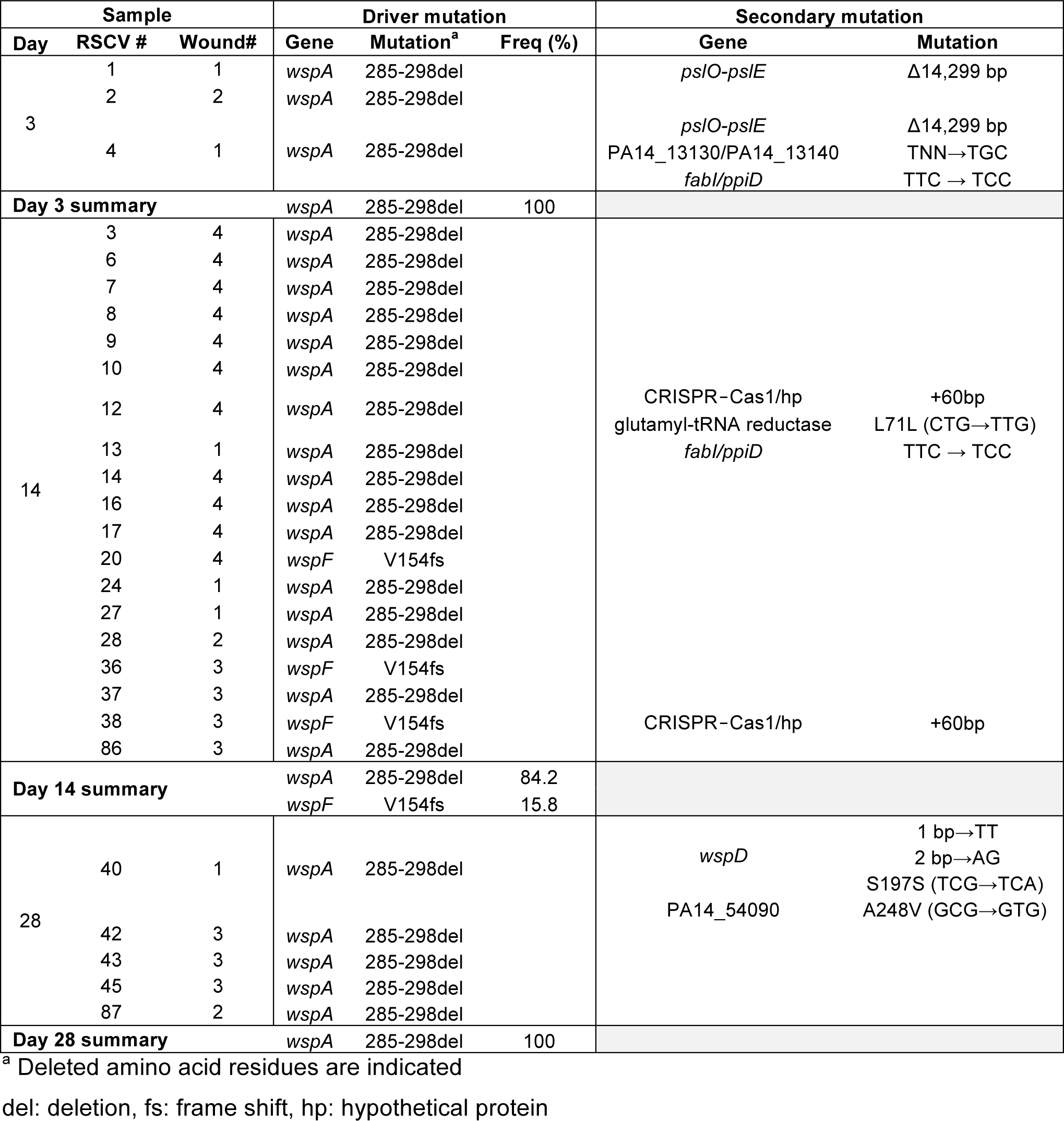
Mutations identified in PA14-1 RSCVs

**Figure 3:**
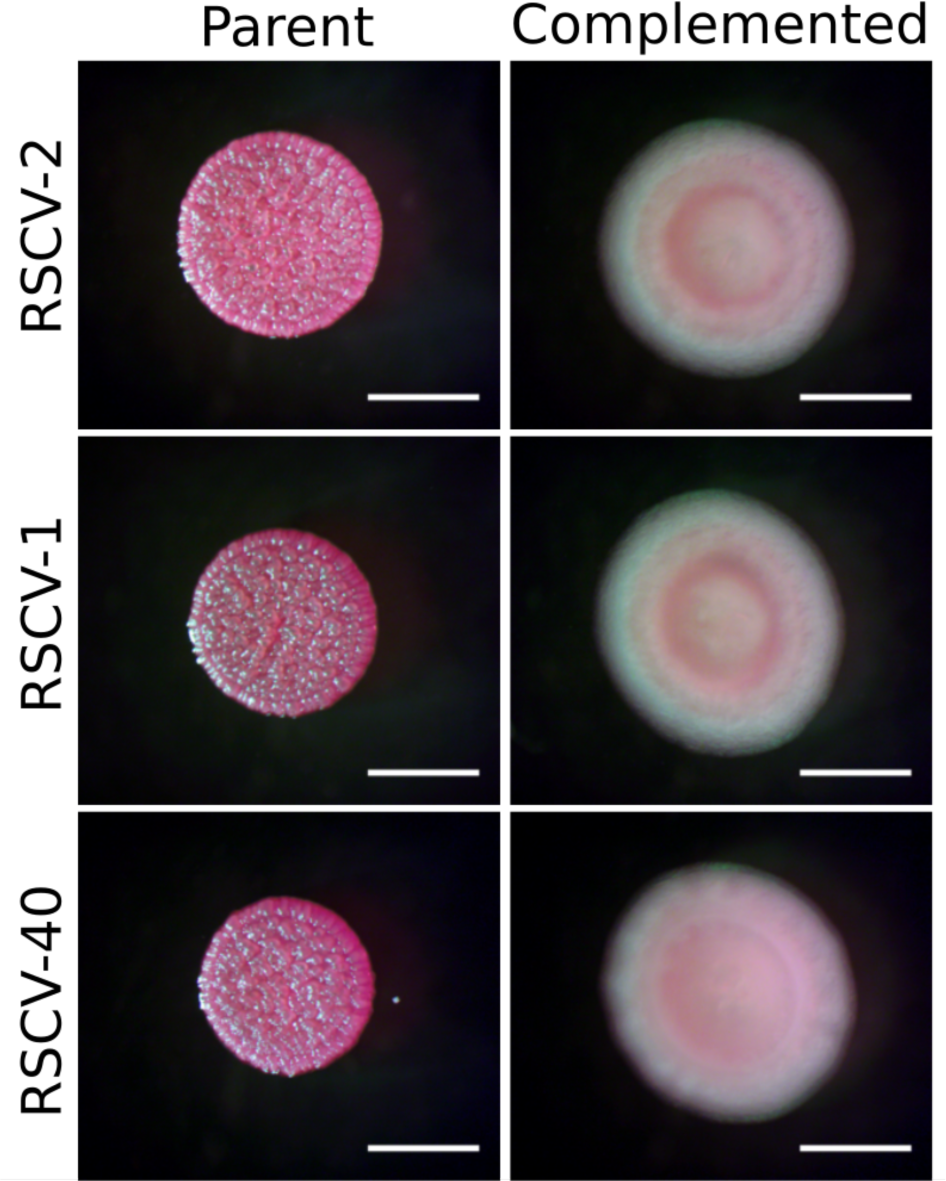
Complementation of the *wspA*Δ42bp mutation. *wspA* was complemented in representative RSCVs by replacing *wspA* Δ285-298 with a wildtype copy on the genome. RSCV-2 was selected as a representative RSCV with the *wspA* driver mutation alone. RSCV-1 and RSCV-40 have the *wspA* driver mutation as well as Δ*pslE*-*pslO* and *wspD* secondary mutations, respectively. Parent and complemented RSCVs (labeled) were grown on VBMM and colony morphology was assessed. Scale bar 2mm.

Some of the RSCVs also possessed secondary mutations (Table 1), demonstrating further evolution in the wound. Two *wspA* RSCVs from 3-d (Table 1; RSCV-1 and RSCV-4) acquired a 14,299bp deletion that removed the remaining *psl* operon. PA14 naturally lacks Psl, since *pslA* – *pslD* are absent^27^. In these two isolates the remaining genes of the *psl* operon, *pslE* – *pslO* were deleted. Both of these PA14-1 RSCVs were isolated from the same wound, however from separate biopsied tissue, suggesting that each deletion may have been a separate event. In the RSCV-1 background, complementation of *wspA* reverted the RSCV colony phenotype to wildtype (Fig 3), indicating that deletion of the remaining *psl* operon did not influence the RSCV phenotype.

Evidence of further evolution of the *wsp* cluster was detected on 28-d. RSCV-40, in addition to having the *wspA* Δ285-298 driver mutation, had 3 separate mutations in *wspD* which led to an early stop codon. This unusual mutation cluster is consistent with error-prone translesion synthesis or DNA template switching facilitated by microhomology^28^. However, as the mutations occur at the *wspD* 3’ end, the WspD N-terminus may still be expressed and functional (Fig S3). WspD is a chaperone, which along with WspB, are predicted to tether WspA and WspE^21^. In this isolate, complementation of *wspA* reverted the RSCV colony phenotype to wildtype (Fig 3), indicating that the *wspD* mutations did not influence the RSCV phenotype. However, this further points towards the strong selective pressure on the Wsp pathway in the chronic infection.

As the driver mutations occurred in the *wsp* pathway, we predicted that the RSCV phenotype was due to overproduction of c-di-GMP. Using both a c-di-GMP *gfp* reporter^29^ and a plasmid encoding the phosphodiesterase PA2133, we determined that both *wspA* and *wspF* mutants had elevated c-di-GMP levels compared to the ancestor PA14-1, and that elevated c-di-GMP levels was responsible for the RSCV colony phenotypes (Supplementary Results; Fig S4). These mutants also showed increased biofilm formation and outcompeted the ancestor strain when grown in *in vitro* planktonic and biofilm competition assays, with greater fitness values seen in the biofilm (Supplementary Results; Fig S5). The presence of secondary mutations did not appear to influence these phenotypes *in vitro*.

We were also interested in identifying how the PA14-1 non-RSCV population adapted to the infection, and if this population acquired mutations that did not result in divergent colony phenotypes. When we sequenced randomly selected PA14-1 non-RSCV isolates, relatively few had acquired chromosomal mutations (Supplemental Results; Table S2). These isolates had similar levels of biofilm formation and metabolic kinetics compared to the ancestor strain (Supplemental Results; Fig S7), suggesting that the RSCVs were the major evolved sub-population within the wounds.

### WspA Δ285-298 mutation leads to auto-induction of the Wsp pathway

As the *wspA* Δ285-298 was the most common driver mutation, we investigated how it may lead to elevated c-di-GMP production. We observed that flanking the junctions of this deletion was a direct repeat at 837-854bp and 921-938bp (Fig S8; bold text^12^).

The WspA Δ285-298 mutation occurs between the predicted HAMP domain (<u>h</u>istidine kinases, <u>a</u>denylate cyclases, <u>m</u>ethyl accepting proteins, and <u>p</u>hosphatases) and the signaling domain (SD; Fig 4A). MCP cytoplasmic domains are comprised of consecutive 7aa heptads^30^. MCPs are defined into classes based on the number of heptads in the cytoplasmic domain^30^, with *P. aeruginosa* WspA belonging to the 40H (40 heptads) MCP class^31^. The 285-298aa (14aa) deletion results in complete deletion of heptads N19 and N16 (Fig 4B).

**Figure 4:**
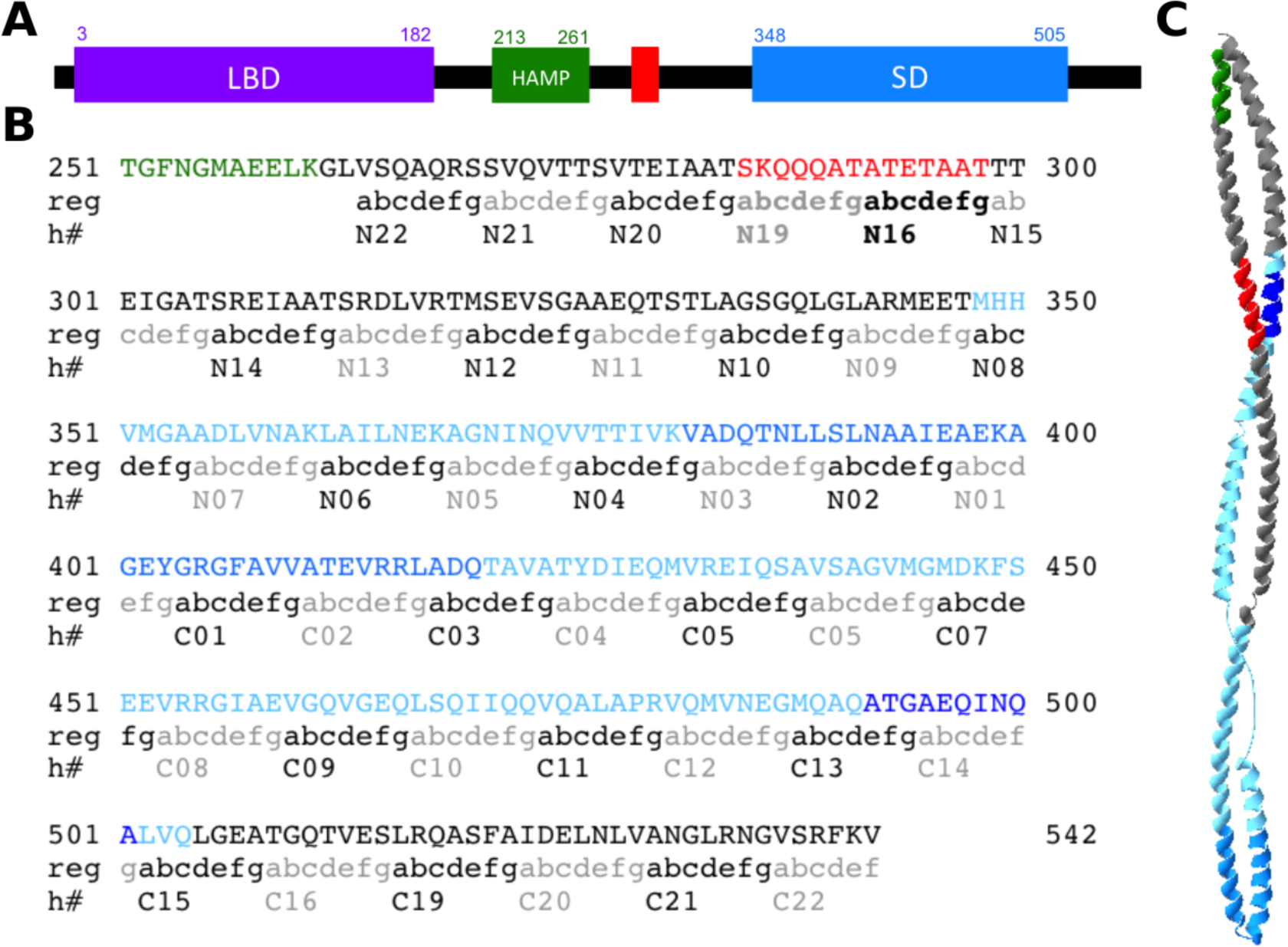
14aa deletion in WspA is predicted to occur opposite the methylation site. (**A)** Schematic of WspA. The different domains of WspA were determined from the Pseudomonas Genome Database^58^ Pfam analysis. LBD = ligand binding domain or the four helix bundle domain (3-182aa). HAMP = linker domain (213-261aa). SD = MCP signaling domain (348-505aa). The region of the 14aa deletion is indicated in red (285-298aa). (**B)** The WspA cytoplasmic domain amino acid sequence. The full amino acid sequence and predicted secondary structure of WspA is depicted in Fig S6. The domains are indicated by the same colors in (**A)**. The signaling domain contains two additional features, the kinase interacting subdomain, or ‘tip’ domain in dark blue (382-420aa) and the predicated methylation site in navy blue (492-501aa)^31^. Both the heptad registers (reg) and the heptad number (h#) are labeled^30^^,^^31^, with consecutive hetpads indicated in alternating black and grey text. (**C)** Homology model of PA14 WspA modeled against the *T. maritime* MCP (PDB 3JA6;^32^) generated using SWISS-MODEL^59^. Colors correspond to the domains indicated in (**B)**. Model spans 250-541aa of WspA.

To observe the localization of the deletion in the protein structure and gain insight into how the Δ285-298 mutation may alter the function and signaling of WspA, we generated a homology model of PA14 WspA against the *Thermotoga maritima* MCP (PDB 3JA6;^32^) (Fig 4C). Our homology model is also supported by the WspA Phyre secondary structure prediction^33^ (Fig S9). Based on the model, the region of the deletion is predicted to occur opposite the methylation site (Fig 4C). We predict that the deletion could de-stabilize or alter the methylation site resulting in auto-induction of WspA and sustained WspR activation.

### Evidence of inter-pseudomonad competition mediated by phages in chronic infections

Two PA14-1 RSCVs, RSCV-12 and RSCV-38, acquired a 60bp insertion at the <u>c</u>lustered <u>r</u>egularly <u>i</u>nterspaced <u>s</u>hort <u>p</u>alindromic <u>r</u>epeat (CRISPR) – <u>C</u>RISPR-<u>as</u>sociated proteins (Cas) locus (Table 1). Both sequences inserted at the intergenic region (−1549/+271) between PA14_33350 (RS13600) and PA14_33370 (RS13605) at the genomic position 2,937,205. The last 28bp of inserted sequence was identical between the two isolates and aligned to the repetitive elements in the PA14 CRISPR array (CRISPR2^34^) (Fig 5A). However, the first 32bp differed, indicative of CRISPR spacer sequences (Fig 5A), which are specific to infective mobile genetic elements^35^. A BLAST search of the inserted CRISPR spacer against the ancestor strains identified that both insertions aligned to different contigs from strain B23-2 (See Supplemental Results; Fig S10).

**Figure 5:**
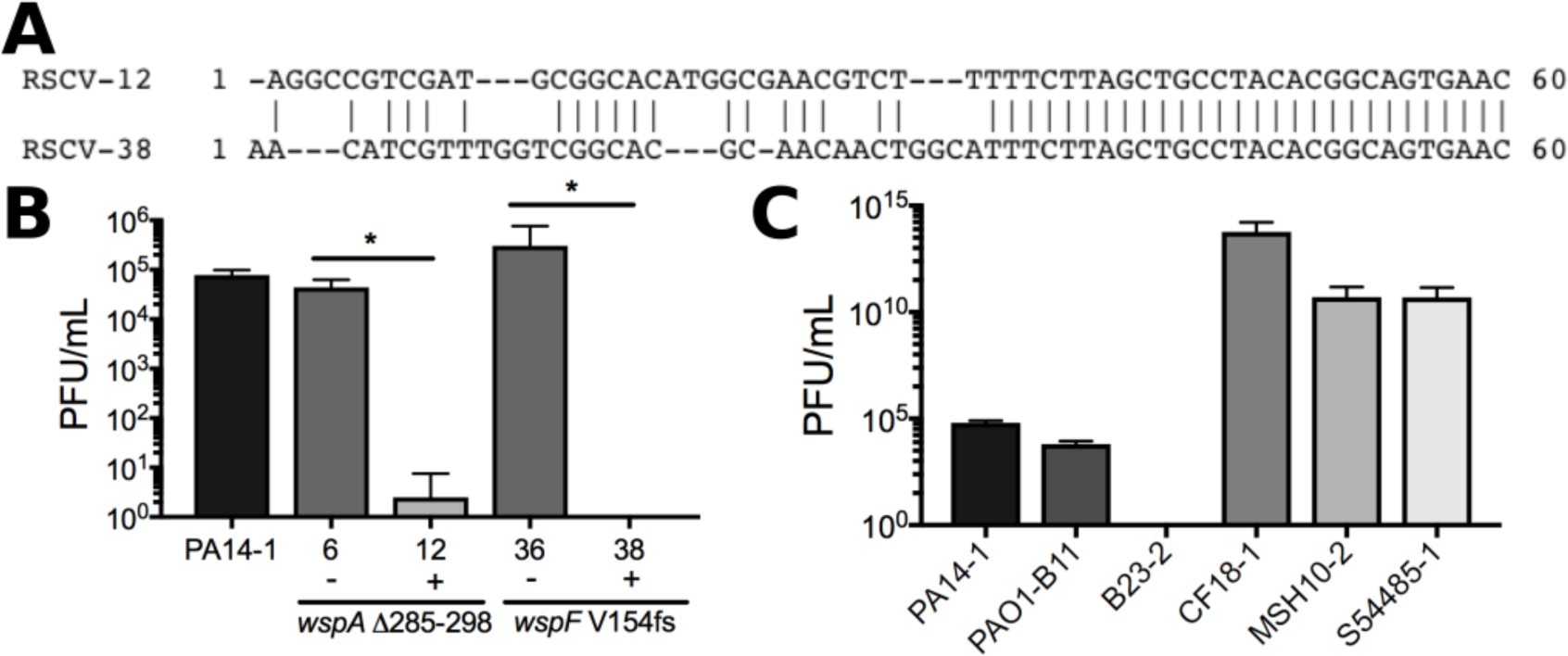
RSCV-12 and RSCV-38 are resistant to infection by phage isolated from B23-2. (**A)** The 60bp insertion sequence in the CRISPR array of RSCV-12 and RSCV-38. Prophages were isolated from B23-2 and plaque assays were performed to determine the level of phage infection for (**B)** representative RSCV isolates and (**C)** the ancestor wildtype *P. aeruginosa* strains. RSCV-6 and RSCV-12 both have the same driver *wspA* mutation, while RSCV-36 and RSCV-38 have the same driver *wspF* mutation. ± indicates the presence/absence of the CRISPR insertion. The driver mutation for each RSCV is labelled. Data presented as mean ± SD, n=4. * p-value <0.05.

We therefore predicted that RSCV-12 and -38 would be resistant to phages isolated from B23-2 due to CRISPR-Cas-acquired adaptive immunity. To test this, we grew B23-2 in mitomycin C and harvested the phage-enriched supernatant. *P. aeruginosa* strains were incubated with the phage lysate and plaque assays were performed. RSCV-12 and RSCV-38 displayed resistance to phage infection. RSCV-38 showed resistance across all replicates (Fig 5B). For RSCV-12, a single plaque was observed in 1 replicate, however for the remaining 3 replicates no plaques were observed (Fig 5B). This indicates that the acquired CRISPR spacers in RSCV-12 and RSCV-38 produced immunity to phages isolated from B23-2. Infection of RSCV-6 and RSCV-36 (CRISPR^−^), which have the same driver mutation as RSCV-12 and RSCV-38 (CRISPR^+^) respectively, revealed that RSCVs were not natively more resistant to phage infection compared to the ancestor strain (Fig 5B). The ancestor strain susceptibility was also determined to assess the host range of the isolated phages. As expected, B23-2 was resistant to phage infection, while the remaining strains showed varying levels of phage sensitivity (CF18-1>MSH10-2, S54485-1>PA14-1, PAO1-B11; Fig 5C).

## Discussion

Here we describe the rapid evolution of adaptive *P. aeruginosa* mutants with conspicuous colony phenotypes arising in a clinically relevant model of chronic infection. There is a consensus in the field that variants arise in an infection as a consequence of adaptation over extended periods of time. However, we isolated RSCVs from early stages of infection, suggesting that variants may evolve more rapidly than originally appreciated. This suggests that RSCVs may be a common, early adaptation during infections and that the selective pressures driving RSCV evolution may persist across these infections.

Even though wounds were infected with six different *P. aeruginosa* strains, we only isolated RSCVs evolved from PA14-1 and PAO1-B11 (Fig 2C). We predict this is due to PA14-1 and PAO1-B11 outcompeting the remaining four strains early in the infection (Fig 1C). *P. aeruginosa* RSCVs were selected in the infection and in *in vitro* conditions (Fig 2C, S5B). The selection coefficients determined here were up to 5 times greater than those identified in the Lenski long-term evolution lines^36^, pointing towards the strong positive selection experienced by RSCVs. Furthermore, *Burkholderia cenocepacia* variants containing *wsp* mutations isolated from an *in vitro* biofilm evolution assay similarly showed high selection coefficients^37^. This suggests that *wsp* mutants experience significant positive selection both *in vivo* and *in vitro* environments. Despite these strong selection coefficients, RSCVs remained at relatively low frequencies in the infection (Fig 2B). This suggests that RSCVs show negative frequency dependence, that is, they exert a strong advantage at low frequencies, but are disadvantageous at high frequencies. Negative frequency dependent selection has been previously observed for evolved rugose variants of *P. fluorescens*^38^^-^^40^. Niche competition^38^^,^^39^ and division of labor^40^ with the ancestor strain drove the evolution of *P. fluorescens* rugose variants from static planktonic and colony growth respectively. In both cases, diversification of the population was maintained by negative frequency dependent selection^38^^-^^40^. The low frequency RSCVs would likely facilitate the ancestor strain in colonizing and establishing biofilms in the wound. We predict that in heterogeneous fitness landscapes, such as those encountered during infection, there are niche environments where the low frequency RSCVs would experience positive dependent frequency selection and become more common. In support of this hypothesis is the observation that the higher RSCV frequencies observed in CF patients are correlated to prolonged exposure to antimicrobials, particularly aerosolized antibiotics^8^^,^^41^.

All the sequenced PA14-1 RSCVs had driver mutations in the *wsp* cluster. This indicates that in chronic wounds the Wsp pathway specifically undergoes selection, and that *wsp* mutants may be more fit than other c-di-GMP-regulating pathways that confer the RSCV phenotype. The *wspA* Δ285-298 was the most common driver mutation and was isolated early in the infection (Table 1). There are two potential explanations for the rapid rise of this single *wspA* mutant in the infection. The first is that it may have been present in the initial inoculum at undetectable levels. The second is that this region may be hyper-mutable owing to the direct repeat (Fig S8). We are currently unable to discern between these two scenarios; however, it is significant that the population rapidly diversifies in the wound due to strong positive selection of adaptive phenotypes provided by *wsp* mutations.

Supporting the second theory is the observation that this region in *wspA* also appears to be under selection in driving RSCV evolution during *in vitro* grown biofilms (Fig S11). The *P. aeruginosa* PAO1 RSCV isolate MJK8, which evolved during biofilm growth in a tube reactor^13^, has an in-frame 66bp deletion (Δ286 – 307aa) in the same region as *wspA* Δ285-298^12^. *P. fluorescens* Pfl01 when grown as a colony biofilm evolved RSCVs with driver mutations identified in *wspC*, *wspA* and *wspE*^40^. One of the *wspA* mutations was a in-frame 84bp deletion (Δ284 – 311aa) again occurring in the homologous region^40^. Finally, *B. cenocepacia* HI2424 RSCVs with *wspA* and *wspE* mutations were isolated from a biofilm bead evolution experiment^37^. While the majority of mutations identified were non-synonymous SNPs, one of the *wspA* mutations was an in-frame 21bp deletion (Δ307-313aa) again in the homologous region^37^ (Fig S11). We predict that these four deletions alter how WspA is methylated/demethylated and ultimately lead to constitutive signaling and auto-induction of the Wsp pathway.

In addition to driver mutations, some PA14-1 RSCVs also gained secondary mutations. Of particular interest was the Δ14,299bp, which deleted the remaining *psl* operon (Table 1; RSCV-1). We predicted that this deletion might lead to increased fitness of the RSCVs over the RSCV driver mutation alone. However, deletion of the remaining *psl* operon did not provide additional fitness benefits under the simple conditions tested. This suggests that in PA14 the remaining *psl* operon (*pslE* – *pslO*) may play a role outside of Psl synthesis, which may have a fitness cost in the wound environment.

Additional secondary mutations of interest were the 60bp insertions in the CRISPR-Cas array of RSCV-12 and RSCV-38 (Table 1, Fig 5A), which encoded resistance to phage(s) isolated from B23-2 (Fig 5C). It has only recently been confirmed that the *P. aeruginosa* type I-F CRISPR-Cas system provides adaptive immunity to phages with a target protospacer^42^. This is dependent on the presence of the correct protospacer adjacent motif (PAM) in the mobile genetic element^42^. In support of this, both protospacers contain the type I-F CRSIPR-Cas specific GG PAM (Fig S10). Furthermore, B23-2 contig-107 contained two additional protospacers to which CRISPR spacers in *P. aeruginosa* have been reported (Table S3, Fig S10A). Of interest was the observation that PA14 already contains a CRISPR spacer identical to a protospacer in contig-107 (Table S3, Fig S10A). This suggests that PA14 had already been exposed to the prophage in B23-2. However, the ancestral PA14-1 was still sensitive to infection (Fig 5B, C), presumably due to the incorrect PAM (Fig S10A). This highlights the importance of insertion of the correct CRISPR spacer in mediating phage immunity. This is only the second report of CRISPR-Cas acquired immunity in *P. aeruginosa* strains^42^ and to our knowledge the first report of CRISPR-Cas adaptive immunity acquired in an infection.

Our data indicate that *P. aeruginosa* experiences strong selective forces in chronic infections, and in response rapidly evolve during the initial stages of infection. RSCVs containing mutations in the *wsp* cluster were the main adapted sub-population isolated from all wounds. This indicates that the Wsp system is the main pathway under selection to evolve adapted variants. This is despite other pathways in c-di-GMP regulation being implicated in RSCV formation, both *in vivo* and *in vitro*^43^^-^^47^. We predict that RSCVs may be an adaptation common to chronic infections and developing therapies that target the RSCV sub-population or prevent their emergence could be transferrable across these infections.

## Materials and Methods

### Bacterial strains, plasmids and media

Bacterial strains and plasmids used in this study are detailed in Table S1. Gene mutant constructs were made using Gibson Assembly (NEB)^48^. Primers used to create the constructs are detailed in Table S4. These were incorporated into the *P. aeruginosa* genome using two-step allelic recombination as previously described^49^. *P. aeruginosa* strains were maintained on LANS (Luria agar no salt; 10g/L tryptone, 5g/L yeast extract solidified with 1.5% agar) unless otherwise specified. RSCV colony morphology was observed on adjusted Vogel-Bonner minimal media (0.2g/L MgSO_4_•7H_2_O, 3.5g/L NaNH_4_HPO_4_•4H_2_O, 10g/L K_2_HPO_4_, 0.1g/L CaCl_2_, 2g/L citric acid, 1g/L casamino acid, 40µg/mL Congo red, 15µg/mL brilliant blue, solidified with 1% agar; VBMM).

For *E. coli* strains, 10µg/mL gentamicin (gent), 100µg/mL ampicillin (amp) or 15µg/mL tetracycline (tet) was used for selection where appropriate. For *P. aeruginosa* strains, 100µg/mL gent, 300µg/mL carbenicillin (carb) or 100µg/mL tet was used for selection where appropriate.

### Porcine full-thickness chronic burn wound model

Swine were housed and studied according to the protocols approved by the Institutional Animal Care and Use Committee (IACUC) at The Ohio State University.

Porcine full-thickness chronic burn wound model was performed as previously described^25^. Briefly, 2 pigs were subjected to thermal injury to achieve 6 full-thickness (third-degree) burns bilaterally and covered with impermeable wound dressings. Burn wounds were infected 3 days post injury with equal amounts of 6 different *P. aeruginosa* strains to achieve a final 250µL inoculum at 10^8^ bacteria (1.6 × 10^7^ each for a total of 1 × 10^8^). The 250µL inoculum was spread over the wounds and allowed to air dry before the wound dressing was re-applied. Wounds were infected with PA14-1, PAO1-B11, B23-2, CF18-1 (GenBank ID; NZ_KI519281), MSH10-2 (GenBank ID; NZ_KE138672) and S54485-1 (GenBank ID; NZ_KI519256). Prior to infection, each strain had been tagged with a unique barcode at the Tn7 site on the genome (see Supplementary Materials and Methods; Table 1).

Wound healing was monitored 3, 14, and 28 days post infection. At each timepoint, 4-8 8mm punch biopsies were taken from 2 wounds on each pig (4 wounds for each timepoint, Fig S2). Biopsies were homogenized in 1mL PBS and plated on *Pseudomonas* isolation agar (PIA) supplemented with 100µg/mL gent for CFUs/ g tissue. To screen for the emergence of adapted *P. aeruginosa* variants, homogenized tissue was also plated onto VBMM supplemented with 100µg/mL gent. Colony morphology variants were passaged onto PIA followed by two rounds on non-selective LANS, before being plated back onto VBMM (without antibiotics) to confirm that the variant phenotype was a result of a stable mutation. Confirmed colony variants were stored at −80°C.

The selection of RSCVs in the wound was determined by calculating the selection coefficient (*s*) according to equation (1)^50^

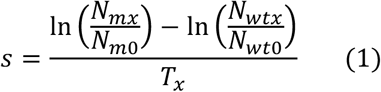

where T_x_ is day x, N is the number of cells, m is the mutant and wt is the wildtype at day x and day 0.

### Colony morphology

1µL of overnight culture was spotted on VBMM plates and incubated at 37°C for 24h. Colonies were imaged on a Stereo Microscope (AmScope) fitted with a Microscope Digital Color CMOS camera (AmScope). Images were processed in FIJI^51^.

### Sequencing and analysis

To determine the frequency of each strain across the infection, genomic DNA was isolated from porcine tissue and the barcodes were sequenced as follows. Tissue was added to 2mL Goodman’s buffer A (100 mM NaCl, 100 mM Tris-HCl pH 8, 10 mM EDTA pH 8, 3.33% SDS, 0.1% sodium deoxycholate) in a bead beater tube with 0.1 mm glass beads and 2-3mm zirconia beads. The tissue was lysed by placing tubes in a TissueLyser II (QIAGEN) for 30s at 50Hz. This was repeated 4 times, placing tubes on ice for 30s in between each lysis round to prevent tubes from overheating. Lysed tissue was incubated with 50µL proteinase K (20mg/mL) for 3h at 55°C. 2mL of phenol, chloroform, isomyl alcohol solution (25:24:1, pH 8) was added and samples were centrifuged for 5min at room temperature. The aqueous phase was removed and mixed with 1.5mL isopropanol. Sample was incubated for 30min at -20°C before being centrifuged for 30min at 4°C. The pellet was washed in 1mL 75% EtOH and again centrifuged. EtOH was removed and the DNA pellet air-dried and resuspended in 500µL sterile water. Strain-specific barcodes were amplified and given Illumina sequencing adapters using Tn7_F and Tn7_R primers indicated in Table S4. The PCR was performed using Expand Long-Template Polymerase (Roche) as described in the Supplementary Materials and Methods. Library sequencing pools were sequenced on NextSeq and MiniSeq High Output SE75 runs at the Petit Institute Molecular Evolution Core Facility at Georgia Institute of Technology. Between 43,789 and 4,938,784 reads were obtained per sample. Analysis script is available on github (https://github.com/glew8/Barcode_Sequencing). Briefly, FastQC 0.11.7 and MultiQC v1.5 were used to confirm sufficient sequencing quality^52^^,^^53^. Cutadapt 1.13 was used to select only those sequences with the insert sequence, isolate barcode sequence, and parse reads containing each barcode^54^. Then egrep was used to count reads with each barcode. As a final check, FastQC was used to identify overrepresented sequences that don’t match any of the queried barcodes.

To identify the ancestor strain that the isolated RSCVs evolved from, colony PCRs were performed using primers specific to each ancestor strain. The forward primer contained the unique barcode used to tag each ancestor strain at the Tn7 site. Therefore, for each RSCV, 6 PCRs were performed. Primers are indicated in Table S4.

To identify mutations, genomic DNA was isolated from colony variants using the DNeasy Blood and Tissue Kit (Qiagen) according to the manufactures protocol. Clonal DNA was sequenced on an Illumina NextSeq 500 using a modified protocol for library prep using the Illumina Nextera kit^55^. 2×151bp sequencing reads for selected isolates were trimmed and quality filtered using Trimmomatic v0.36 (settings: LEADING:20 TRAILING:20 SLIDINGWINDOW:4:20 MINLEN:70)^56^. The reads passing quality filtering were then used for variant calling with the open-source program *breseq* v0.30.0 using default settings^57^. The reference sequences for variant calling were acquired from NCBI‘s RefSeq database (NC_002516.2 for PAO1, NC_008463.1 for PA14).

### Homology modelling

The PA14 WspA sequence was obtained from the Pseudomonas genome database^58^. The sequence was submitted to SWISS-MODEL using a template search^59^. Quality of the returned homology models was assessed on the Global Model Quality Estimation (GMQE) score (numbers closer to 1 indicate more accurate model) and the QMEAN score (numbers closer to 0 indicate that the model is comparable to experimental structures). The homology model of PA14 WspA against the MCP of *Thermotoga maritima* (PDB 3JA6;^32^) was determined to be the most accurate. The homology model had a GMQE score of 0.37 and QMEAN score of -1.68.

### Prophage isolation and plaque assay

To isolate prophages from B23-2, an overnight culture of B23-2 was diluted 1:100 and incubated for 30min at 37°C shaking at 200rpm. 0.5µg/mL of mitomycin C was added to the culture and the OD_600nm_ was measured. The culture was re-incubated and the OD_600nm_ measured every h. When the OD_600nm_ began to decrease the cells were pelleted by centrifugation and the supernatant filter sterilized and stored at 4°C.

To determine the level of susceptibility of *P. aeruginosa* strains to the bacteriophage(s) isolated from B23-2, 100 – 200µL of mid-log *P. aeruginosa* culture was incubated with 100µL serial dilutions of bacteriophage lysate for 15min at 37°C. The infection was added to 5mL molten soft agar (LB solidified with 0.7% agar) supplemented with 10mM CaCl_2_ and MgSO_4_. This was then poured over solidified hard agar (LB solidified with 1.5% agar), allowed to solidify and incubated overnight. The number of resulting plaques was then counted and PFU/mL determined.

### Statistical Analysis

Data are presented as mean ± SD. To determine if data conformed to a normal distribution a Shapiro-Wilk test was performed. All of the data sets were normally distributed except for the PFU/mL data (Figure 5). For these data sets, means were compared using the non-parametric t-test. All other comparisons were made using a one-way ANOVA with a Tukey’s post-hoc test and Student’s t-test. Analyses were performed using GraphPad Prism v.5 (Graphpad Software). Statistical significance was determined using a p-value <0.05.

### Data Availability

All sequencing reads from isolates are deposited in NCBI SRA under Bioproject number PRJNA491911 and Biosample accession numbers SAMN10101410 - SAMN10101459.

## Acknowledgements.

We would like to thank Michael Kann for his help with the planktonic and biofilm fitness assays.

